# AmbiDose: per-cell removal budgets for ambient RNA contamination

**DOI:** 10.64898/2026.09.27.754849

**Authors:** Zhao Li, Aaron W. James, Shengxuan Li

**Affiliations:** Department of Pathology, Johns Hopkins University, Baltimore, MD 21205, USA

## Abstract

Ambient RNA contamination confounds expression analysis in droplet-based single-cell and single-nucleus RNA sequencing. Here we present AmbiDose, which combines an ambient profile estimated from empty droplets with cell-specific nominal removal budgets and expression-based protection of lineage-enriched genes. Its ambient dose guides count removal rather than estimating a calibrated contamination fraction. In a held-out human–mouse GEM-X mixture, AmbiDose removed 92.0% of cross-species UMIs while retaining 99.6% of same-species UMIs. It reduced off-target marker expression in kidney-tumor and fetal-liver cohorts and yielded the lowest proximal-tubule marker leakage among the tested methods across five strain-mixed kidney libraries. Simulations supported the contribution of cell-specific dose estimates but identified reduced endogenous retention or incomplete removal when ambient and endogenous expression overlapped. AmbiDose is available as a Python package compatible with Scanpy.

---

Droplet-based single-cell and single-nucleus RNA sequencing (scRNA-seq and snRNA-seq) quantify gene expression in individual cells or nuclei. RNA released by stressed or lysed cells can be co-encapsulated and counted alongside endogenous transcripts. This ambient RNA produces counts in empty droplets and off-target expression in cell-containing droplets, reducing marker specificity and confounding cell annotation and differential expression analysis ^1–5^.

Existing methods differ in how they estimate ambient composition and distinguish background from endogenous expression. SoupX estimates the ambient profile from empty droplets and, with clusters, uses autoEstCont to choose informative genes and to estimate a contamination fraction (default prior priorRho = 0.05) before cell-level adjustCounts1. DecontX models each cell as a mixture of endogenous and contaminating expression, with an optional empty-droplet background ^2^. CellBender jointly models cell-containing and empty droplets to infer background counts ^3^. scCDC identifies contamination-associated genes across clusters and subtracts a fixed count per gene within affected clusters ^6^. Other approaches model ambient noise or target selected genes ^7–9^.

A shared ambient composition does not imply equal contamination across droplets. Genotype-resolved kidney data show substantial variation in background levels among cells within a library ^10^. Although DecontX and CellBender already estimate cell-specific contamination, separating ambient from endogenous expression remains difficult when abundant lineage transcripts dominate the background or when both sources express the same genes ^2,3,11^. These observations motivate an approach that links cell-specific evidence of contamination to count removal while using cell-type expression structure to protect lineage-enriched transcripts.

Here we introduce AmbiDose, which combines an empty-droplet ambient profile with cell-specific nominal removal budgets and expression-based gene protection (Fig. 1). The dose is an operational score, not a calibrated contamination fraction. Cell-type mean expression identifies genes used to estimate the dose and genes protected during subtraction. We evaluated the method using species mixtures, primary tissues, controlled contamination simulations and strain-mixed kidney libraries. A GEM-X mixture and a dorsal-root ganglion library were held out from parameter and feature selection.

**Fig. 1.**
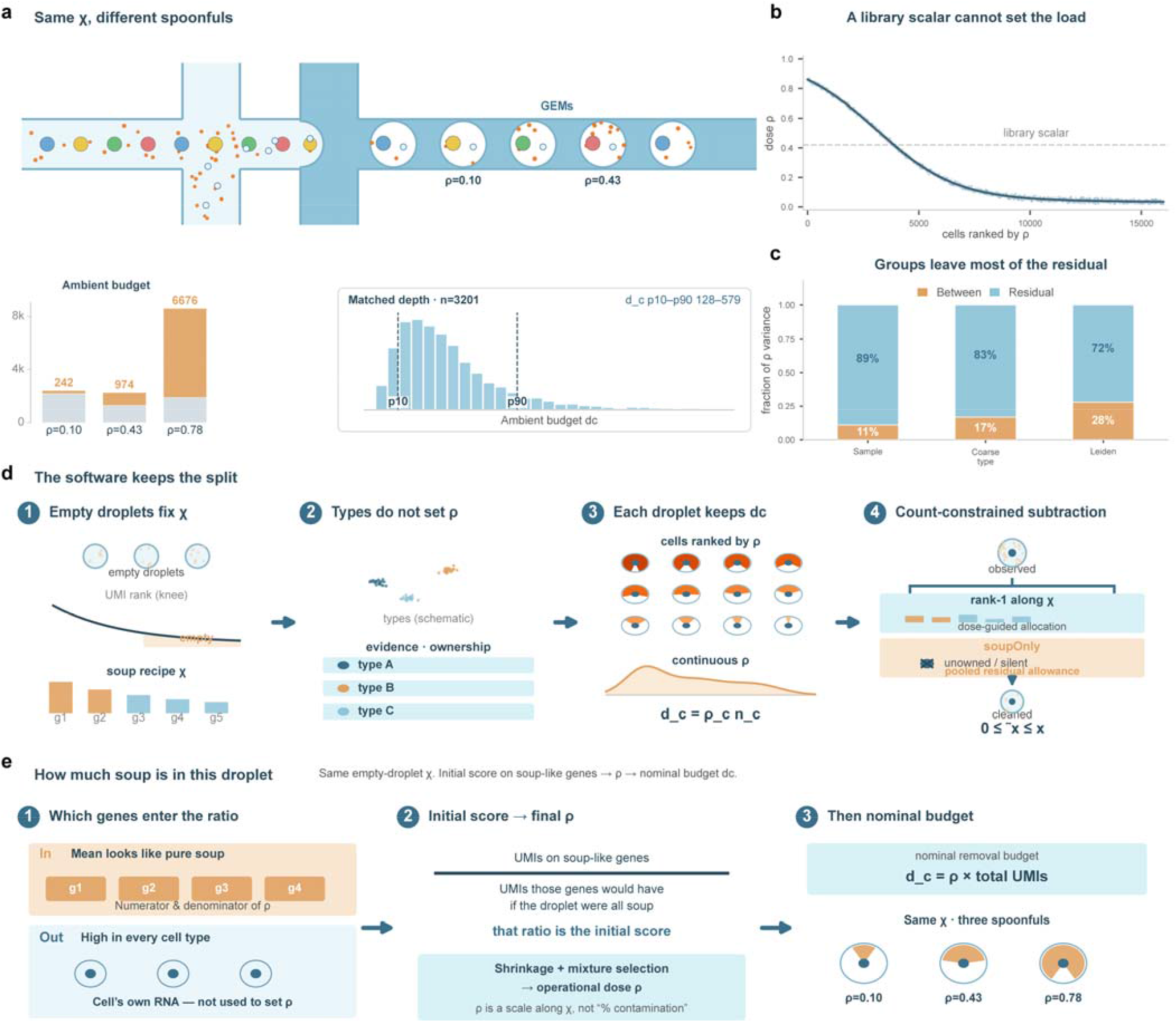
Cell-specific dose estimates under a shared ambient profile. (a) Shared empty-droplet profile χ and example droplets. Three T cells from one kidney-tumor library have ρ = 0.10, 0.43 and 0.78; bars show nominal removal budgets d_c_ of 242, 974 and 6,676 UMIs. The histogram shows d_c_ for 3,201 cells in a narrow UMI band. (b) Cells ranked by estimated ρ; the dashed line is a library-wide reference value. (c) Between-group and within-group fractions of variance in estimated ρ for sample, coarse type and Leiden cluster. (d) Workflow: empty droplets estimate χ; type-level expression identifies evidence and protected genes; cell-specific doses guide count allocation. The released implementation guarantees 0 ≤ corrected count ≤ raw count, but does not impose a strict per-cell ceiling of d_c_. (e) Initial score based on the ratio of observed evidence-gene counts to their expected counts under a purely ambient profile. Shrinkage and mixture-based selection produce the final operational dose (Methods; Supplementary Note 1).

## Results

### Overview of AmbiDose

AmbiDose first estimates a sample-specific ambient profile, χ, from pooled empty-droplet counts (Fig. 1d,e). For each cell, it calculates an initial score from genes whose type-level expression is compatible with the ambient profile, applies shrinkage and selects between this estimate and a fixed-ambient-profile mixture estimate (Methods). The resulting dose, ρ_c_, defines a nominal removal budget, d_c_ = ρ_c_ n_c_, where n_c_ is the total UMI count. Coarse cell types, supplied by the user or inferred by Leiden clustering, determine evidence and protected-gene sets. Subtraction combines profile-directed removal with additional clearance of genes classified as unexpressed and unprotected within a type. Budgets guide this allocation but are not strict per-cell removal ceilings in the released implementation. Corrected counts remain nonnegative and do not exceed the corresponding input counts.

### Estimated ambient dose varies within cell groups

Cells sharing an ambient profile can have different estimated doses. Three T cells from one kidney-tumor library had ρ values of 0.10, 0.43 and 0.78 and nominal budgets of 242, 974 and 6,676 UMIs (Fig. 1a). Across the analyzed cells, sample, coarse type and Leiden cluster explained 11%, 17% and 28% of the variance in estimated ρ, respectively (Fig. 1b,c). These estimates describe model output rather than independently measured contamination. Consistent with cell-to-cell heterogeneity, cross-species UMI counts varied 1.3–1.7-fold between the 10th and 90th percentiles within species and a narrow library-size window; estimated ρ varied 1.3–1.6-fold in the same windows (Supplementary Fig. 1). The contribution of cell-specific dose estimates is tested by ablation below (Fig. 4f).

### Validation in human–mouse mixtures

We used cross-species UMIs in confidently assigned singlets as an observable proxy for contamination. Sensitivity measures cross-species UMI removal, specificity measures same-species UMI retention, and precision is the fraction of removed UMIs that are cross-species (Fig. 2). Same-species retention does not measure endogenous retention because same-species ambient RNA is also included. AmbiDose achieved sensitivity, specificity and precision of 0.954, 0.997 and 0.943 on hgmm12k, and 0.940, 0.999 and 0.950 on Mixture. Under the tested configuration, CellBender reached sensitivity 0.2353, specificity 0.99925 and precision 0.9435 on hgmm12k, and 0.0890, 0.99863 and 0.4608 on Mixture: it retained nearly all same-species UMIs but removed a smaller fraction of cross-species UMIs.

**Fig. 2.**
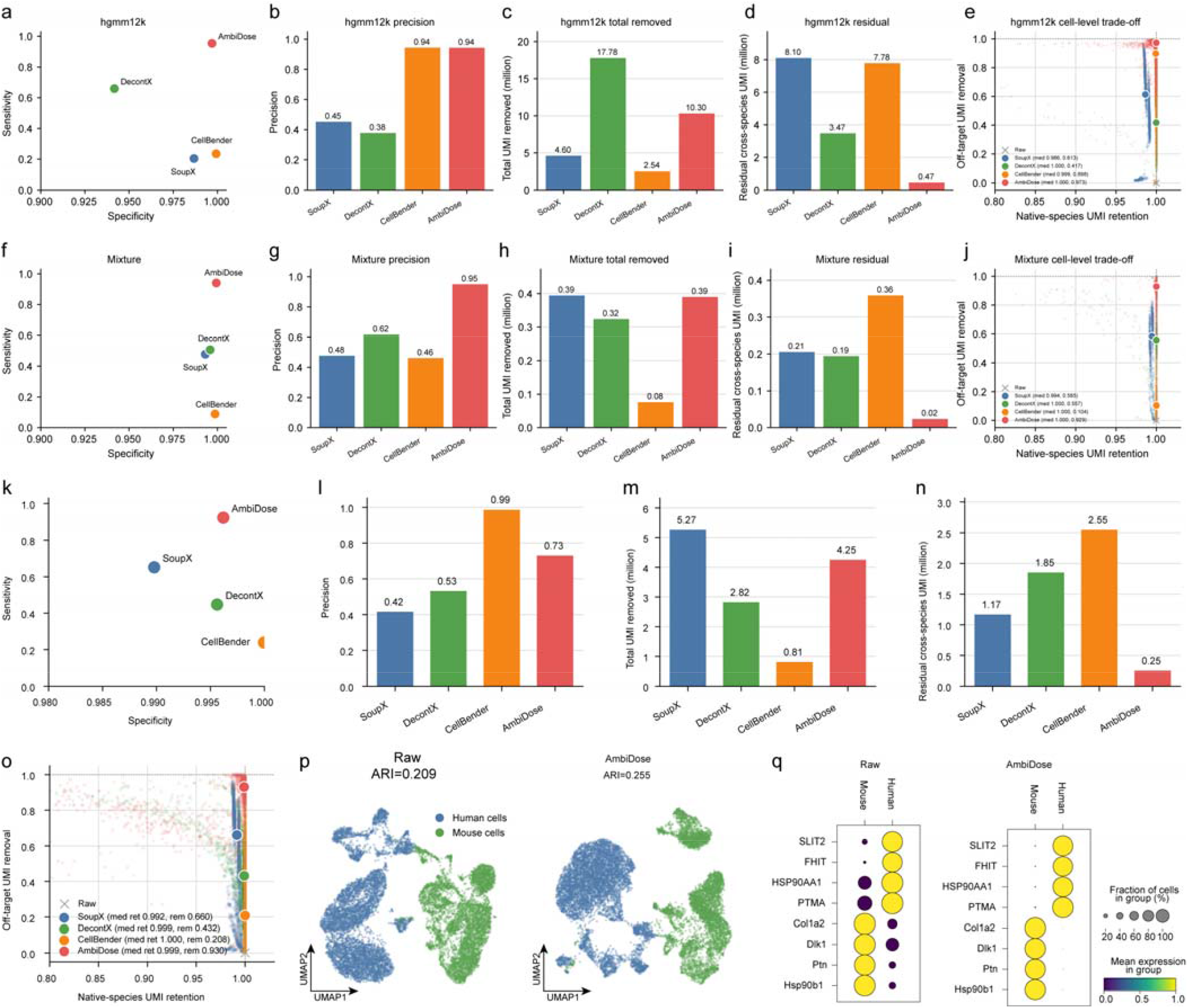
Species mixing on hgmm12k. (a–e), Mixture (f–j) and the held-out GEM-X library (k–q). SoupX, blue; DecontX, green; CellBender, orange; AmbiDose, red; raw, gray. Comparator configurations are given in Methods and Supplementary Table 3; a DecontX rerun with empty-droplet background is given in Supplementary Table 7. (a, f, k) Sensitivity (fraction of cross-species UMIs removed) against specificity (fraction of same-species UMIs retained). (b, g, l) Precision of the removed UMIs. (c, h, m) Total UMIs removed (millions). (d, i, n) Cross-species UMIs remaining (millions). (e, j, o) Per-cell same-species UMI retention against off-target UMI removal for cells with at least one raw off-target UMI; large points are medians. (p) Raw and AmbiDose UMAPs of the GEM-X mixture colored by species (raw human versus mouse UMI dominance); ARI compares these labels with Leiden clusters on the same matrix. (q) Species-marker dot plots; dot size, fraction of cells with a detected count; color, mean log1p expression.

On GSE147203 ^10^, a spike-in-validated dataset included in method development, AmbiDose achieved sensitivity 0.956, specificity 0.996 and precision 0.844. CellBender reached sensitivity 0.421, specificity 0.99994 and precision 0.993; SoupX and DecontX reached sensitivities of 0.447 and 0.758. DecontX used no empty-droplet background in the main comparisons; supplying that background increased its sensitivity on all four species-mixture libraries, with changes in same-species retention and precision (Supplementary Table 7).

We next applied the locked settings to an independent GEM-X 3′ human–mouse mixture. Among 11,875 confidently assigned singlets, AmbiDose removed 92.0% of cross-species UMIs, retained 99.6% of same-species UMIs and achieved precision 0.730, leaving 0.25 million cross-species UMIs (Fig. 2k–n). Median per-cell same-species retention and cross-species removal were 0.999 and 0.930 for AmbiDose, 0.992 and 0.660 for SoupX, 0.999 and 0.432 for DecontX, and 1.000 and 0.208 for CellBender (Fig. 2o). Both species separated in the raw UMAP ^12^; agreement between Leiden clusters and species labels increased from ARI 0.209 to 0.255 after AmbiDose correction ^13^ (Fig. 2p,q). scCDC achieved sensitivity 0.135, specificity 0.977 and precision 0.063 under its tested defaults (Supplementary Table 2).

### Off-target marker expression in primary tissues

In primary tissues, we quantified off-target marker expression using the leak ratio, defined as the mean expression of other types’ markers divided by the mean expression of a type’s own markers (Methods). Among 16,282 matched kidney-tumor cells ^14^, AmbiDose reduced the ratio from 1.07% to 0.83% while retaining 94.4% of on-target marker UMIs; the ratios after SoupX, DecontX and CellBender correction were 0.97%, 0.84% and 1.00%. In the 39-sample fetal-liver cohort ^15^ (114,308 matched cells), AmbiDose reduced the ratio from 3.94% to 1.30% while retaining 85.5% of on-target marker UMIs, compared with ratios of 2.19%, 2.69% and 3.27% for the three comparators (Fig. 3a).

**Fig. 3.**
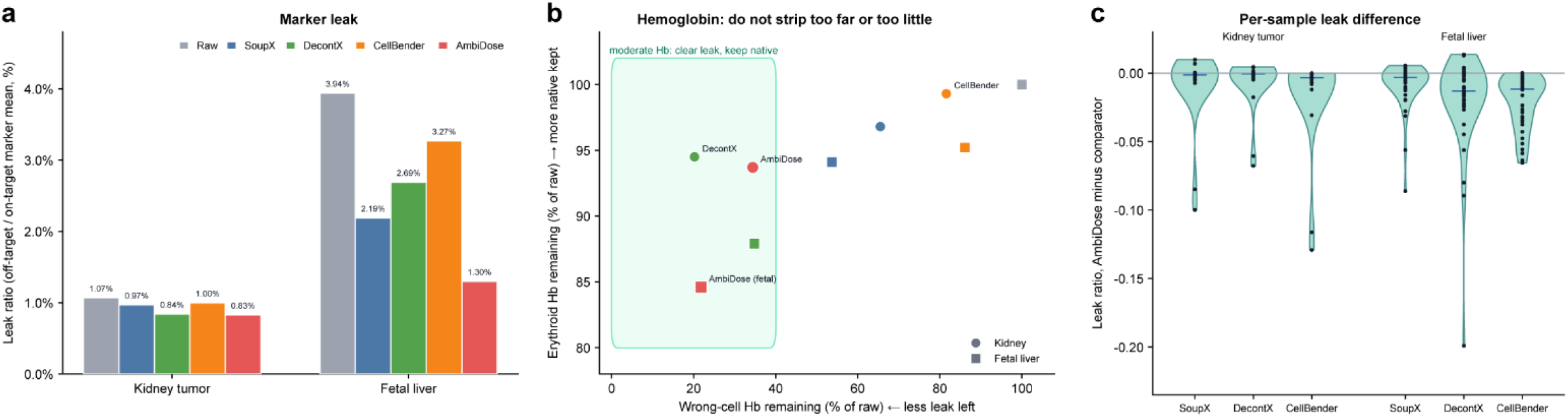
Off-target marker expression and hemoglobin retention in matched cells. (a) Leak ratio, defined as mean off-target marker expression divided by mean on-target marker expression, in kidney-tumor and fetal-liver cohorts. (b) Hemoglobin remaining in non-erythroid versus erythroid cells, each expressed as a percentage of raw expression. Circles, kidney tumor; squares, fetal liver. The shaded box indicates the displayed reference region for off-target clearance and erythroid retention. (c) Library-level differences in leak ratio between AmbiDose and each comparator (kidney tumor, n = 23 libraries; fetal liver, n = 39). Negative values favor AmbiDose. Donor-level comparisons are reported in the text.

Because libraries from the same donor are not independent, donor-level comparisons provide the more relevant biological inference. After averaging libraries within donor, AmbiDose had lower leak ratios than CellBender and DecontX in all 14 fetal-liver donors (both P = 1.2 × 10^−4^), and lower ratios than SoupX in 11 of 14 donors (P = 0.0052). In kidney, ratios were lower than CellBender in all seven donors (P = 0.016, unadjusted), but differences from SoupX and DecontX were not significant (P = 0.11 and 0.47). Library-level comparisons are shown in Fig. 3c and Supplementary Tables 5 and 6. Of the kidney comparisons, only the difference from CellBender met the library-level Bonferroni threshold of 0.0083; all three fetal-liver comparisons met this threshold.

Hemoglobin expression provided a complementary measure of the balance between off-target removal and on-target retention (Fig. 3b). In kidney tumor, AmbiDose retained 93.7% of erythroid hemoglobin and 34.5% of non-erythroid hemoglobin; DecontX retained 20.2% of non-erythroid hemoglobin. In fetal liver, erythroid hemoglobin retention was 84.6% for AmbiDose and 87.9% for DecontX. Thus, AmbiDose did not achieve the strongest off-target clearance or on-target retention in every comparison.

### Component ablation and limits under simulated contamination

We used two contamination-injection designs to examine component contributions and failure modes. These simulations provide controlled tests rather than estimates of typical tissue performance.

In PBMC4k ^16^, a six-gene injection panel comprised genes absent from the clean background (INS, GCG, CPA1, ALB, MYH6 and TNNT2). In the zero-injection control, AmbiDose retained 99.999% of observed UMIs. At 2% library contamination drawn along the empty-droplet profile, it removed 66% of injected UMIs at entries with zero endogenous counts and retained 99.1% of endogenous UMIs. At 5% contamination, the corresponding values were 60% and 97.6%. The sparse-panel evaluation under default settings also yielded a reported mean recovery of 17.7% (gene range, 10.1–24.5%). Recovery was incomplete in this evaluation.

The second design followed the high-ambient cell-line mixture of Cargnelli et al. ^4^: an equal mixture of three breast cell lines, with cell-specific ambient fractions drawn from N(0.6, 0.3) and clipped to [0.01, 0.8] (Fig. 4). The lines share many expressed genes, making ambient and endogenous contributions difficult to distinguish. We therefore report bounds on the ambient removal score (ARS) and endogenous retention score (ERS), reflecting the possible attribution of removed counts at entries containing both sources (Methods).

**Fig. 4.**
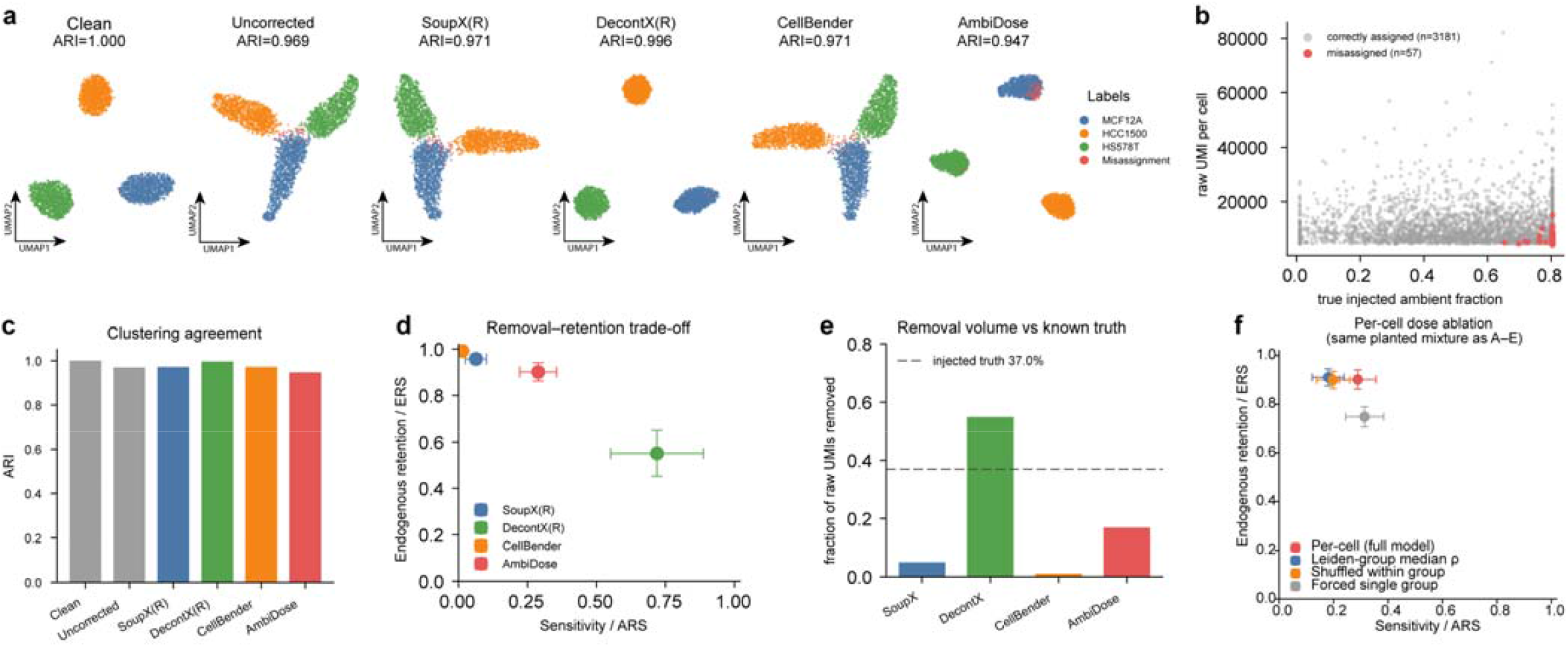
Component ablation and a high-ambient simulation. Equal mixture of MCF12A, HCC1500 and HS578T cells, with cell-specific ambient fractions drawn from N(0.6, 0.3) and clipped to [0.01, 0.8]. (a) UMAPs of the clean, uncorrected and corrected matrices, colored by cell line; misassigned cells are red. (b) Raw UMI counts versus simulated ambient fraction for AmbiDose; gray, correctly assigned cells (n = 3,181); red, misassigned cells (n = 57). (c) ARI with cell-line identity. (d) ARS and ERS bounds obtained by assigning removal at mixed-source entries preferentially to ambient or endogenous counts. Bars show attribution bounds, not confidence intervals. (e) Fraction of raw UMIs removed; the dashed line marks the library-wide injected fraction of 37.0%. (f) Full model, Leiden-group median dose, within-group shuffled dose and single-group ablations. The first two dose substitutions keep χ, ownership and the subtraction procedure fixed; the single-group analysis also changes type structure. Bars are defined as in d.

AmbiDose achieved ARS 0.222–0.356 and ERS 0.862–0.941 (Fig. 4d). DecontX without an empty-droplet background removed more injected ambient RNA but retained less endogenous RNA. SoupX and CellBender removed smaller fractions of the injected ambient RNA; CellBender achieved ARS approximately 0.000–0.001 and ERS approximately 1.000 under the tested configuration. Total UMI removal was 16.9% for AmbiDose, 54.9% for DecontX, 5.0% for SoupX and 1.05% for CellBender, compared with a library-wide injected ambient fraction of 37.0% (Fig. 4e).

Correction did not improve cell-line clustering in every case. AmbiDose yielded ARI 0.947, compared with 0.969 for the uncorrected matrix, 0.971 for SoupX and CellBender, and 0.996 for DecontX (Fig. 4a,c). The 57 misassigned cells had higher simulated contamination and lower raw UMI counts than the 3,181 correctly assigned cells (Fig. 4b), consistent with greater vulnerability of low-count cells to signal loss during correction.

The same simulation tested the contribution of cell-specific dose estimates. The nominal budget d_c_ correlated with the intended ambient count (Spearman 0.96 overall and 0.91–0.95 within library-size quintiles), but the regression slope was 0.29, indicating substantial underestimation on this design (Supplementary Fig. 2). With χ, ownership and the subtraction procedure fixed, replacing each cell’s dose with its Leiden-group median reduced ARS to 0.121–0.239 (ERS 0.876–0.945); shuffling doses within groups reduced ARS to 0.137–0.259 (Fig. 4f). Assigning all cells to one group before dose estimation and subtraction increased ARS to 0.244–0.384 but reduced ERS to 0.707–0.789. These comparisons support contributions from both cell-specific dose information and type structure under the tested simulation.

### Marker specificity in strain-mixed kidney libraries

GSE218853 contains five mouse kidney libraries generated by pooling cells from two subspecies, allowing cross-genotype signal to be identified at informative SNPs ^11^. Three libraries contain single cells (rep1–rep3) and two contain nuclei (nuc2 and nuc3). We included nuc3, which the original study excluded from downstream analyses because of poor performance across the methods tested. Informative-SNP coverage was lower in nuc2 (mean, 127 per cell) than in the other libraries (564–816). Raw proximal-tubule (PT) marker leakage was 25.0% in nuc2 and 23.9% in nuc3 (Fig. 5).

**Fig. 5.**
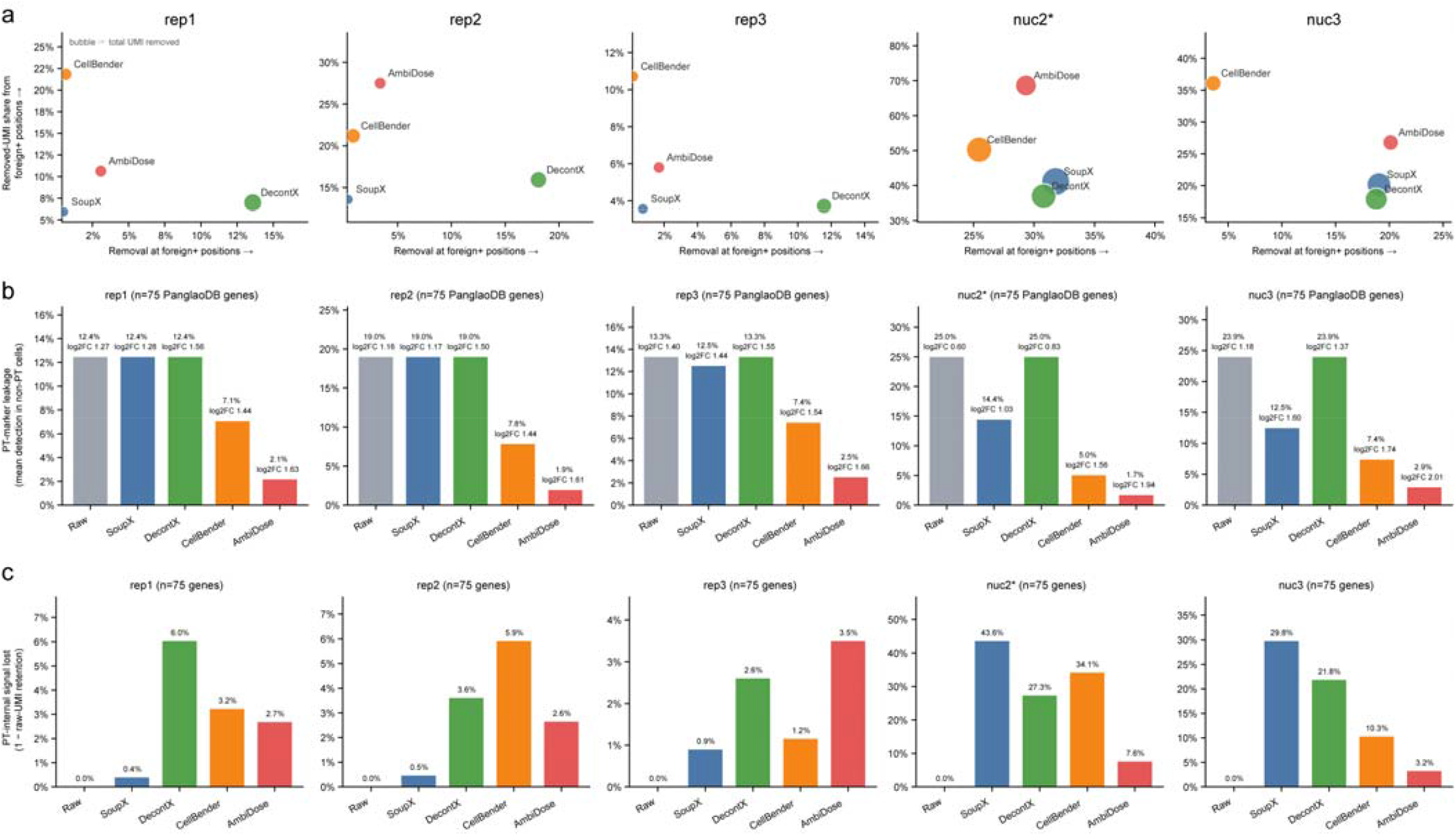
Marker and genotype-position readouts in GSE218853 kidney libraries. Single-cell (rep1–rep3) and single-nucleus (nuc2, nuc3) libraries; nuc2* has 4–6-fold fewer informative SNPs per cell than the other libraries. SoupX, blue; DecontX, green; CellBender, orange; AmbiDose, red. (a) Mean removal at foreign-positive positions against the share of removed UMIs that came from those positions; bubble area is proportional to total UMIs removed. (b) PT-marker leakage (mean detection in non-PT cells) and log2 fold change for the 75 PanglaoDB proximal-tubule markers present in the data. (c) PT-internal signal lost (1 − raw-UMI retention) for the same genes within PT cells.

Using the 75 PanglaoDB PT markers present in these data ^17^, AmbiDose yielded the lowest mean detection in non-PT cells across all five libraries (1.7–2.9%, versus 5.0–7.8% for CellBender and 12.4–25.0% for SoupX and DecontX) and the highest PT-versus-rest log2 fold changes (1.61–2.01, versus 0.60–1.40 on the raw matrices; Fig. 5b). On the two nucleus libraries, it also had the smallest loss of PT-internal marker UMIs: 7.6% in nuc2 and 3.2% in nuc3, compared with 43.6% and 29.8% for SoupX, 27.3% and 21.8% for DecontX, and 34.1% and 10.3% for CellBender (Fig. 5c). On the single-cell libraries, AmbiDose lost 2.6–3.5% of PT-internal signal, more than SoupX on all three libraries and more than CellBender on rep3. These results support improved marker specificity with a favorable retention trade-off in the two high-background nucleus libraries under the tested configurations.

Genotype information provided a distinct, position-level readout (Fig. 5a). For AmbiDose, mean removal at foreign-positive positions ranged from 0.017 to 0.293, and retention at foreign-negative positions ranged from 0.882 to 0.996. AmbiDose was on the Pareto front for nuc2 and nuc3 when removal at foreign-positive positions was compared with the share of removed UMIs originating from those positions. On rep1, rep3 and nuc3, CellBender combined the highest removed-UMI share from foreign-positive positions with low removal at those positions. Because each position can contain both foreign and endogenous molecules, these scores do not identify the molecular origin of the removed counts.

Total UMI removal differed most on the nucleus libraries. In nuc2, AmbiDose removed 18.0% of UMIs, compared with 35.6% for SoupX, 27.2% for DecontX and 28.4% for CellBender. scCDC achieved mean removal at foreign-positive positions of at most 0.017 across the five libraries under its tested defaults (Supplementary Table 2).

### Application to a held-out dorsal root ganglion library

We applied the locked settings to an uninjured mouse dorsal root ganglion library (GSE265817, GSM8229618) ^18^, which was not used for parameter or feature selection. Major neuronal and non-neuronal groups were visible in independently clustered raw and corrected matrices (Fig. 6a,b; Leiden resolution, 0.5); the corrected map omits 42 unassigned cells. After correction, Fabp7, Apoe and Mpz showed more restricted expression across the annotated classes, with Apoe retained in immune cells, and Tubb3 expression was concentrated in neurons (Fig. 6c–h).

**Fig. 6.**
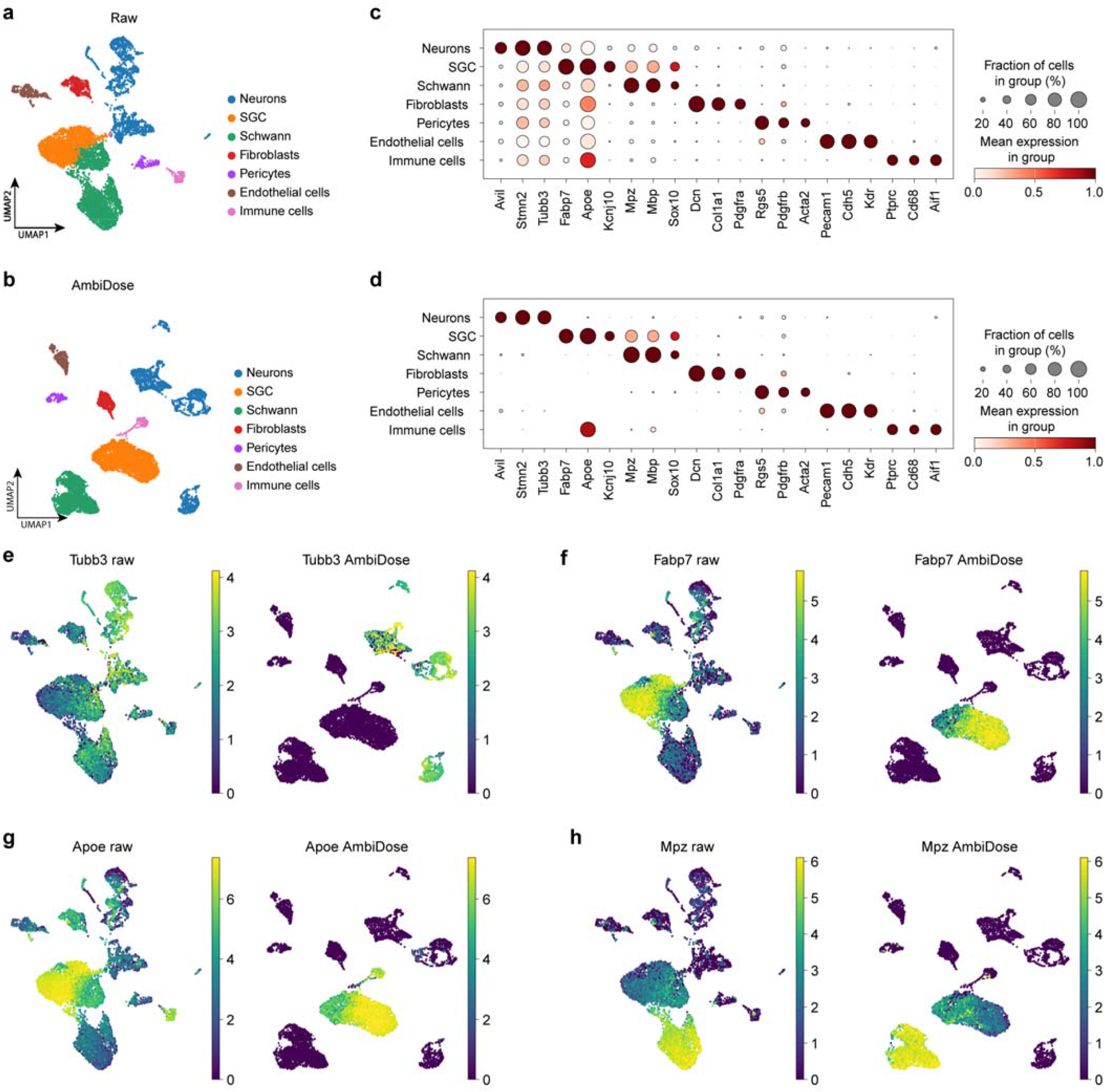
Held-out mouse dorsal root ganglion (GSE265817, GSM8229618). (a, b) UMAPs of the raw and AmbiDose matrices, each clustered independently (Scanpy; Leiden resolution 0.5); colors are cluster labels. The AmbiDose panel omits 42 unassigned cells. (c, d) Marker dot plots, three genes per class; dot size, fraction of cells in the class with a detected count; color, gene-scaled mean. (e–h) Raw and AmbiDose expression of Tubb3, Fabp7, Apoe and Mpz; each pair shares one color scale.

Collectively, these analyses show that an empty-droplet ambient profile combined with per-cell nominal doses and expression-based protection can reduce identifiable contamination while retaining substantial endogenous signal across the droplet and nuclear libraries tested here. Against SoupX, DecontX, CellBender and scCDC under the reported configurations, AmbiDose improved the removal–retention balance in several settings, including foreign-allele and proximal-tubule marker readouts in strain-mixed kidney nuclei. The held-out GEM-X mixture and dorsal root ganglion library indicate that the same locked settings transfer beyond the datasets used for method development.

## Discussion

AmbiDose combines an ambient profile estimated from empty droplets, cell-specific nominal removal budgets and expression-based protection of lineage-enriched genes. Across the tested datasets, it removed cross-species signal and reduced off-target marker expression while retaining substantial on-target signal. The held-out GEM-X mixture and dorsal root ganglion library extend evaluation beyond the datasets used for method development.

The methodological contribution is the combination of cell-specific dose information with type-aware count allocation. Per-cell estimation itself is not unique to AmbiDose: DecontX and CellBender also model cell-specific contamination ^2,3^. Our ablations show that replacing individual doses with group medians or shuffled doses reduced ambient removal in the tested simulation, whereas removing type structure reduced endogenous retention. These results support the use of both components but do not establish calibrated contamination estimates or a universal advantage across datasets.

Comparator performance depended on configuration and dataset. DecontX used no empty-droplet background in the main comparisons; supplying that background changed its species-mixture results (Supplementary Table 7), consistent with the importance of background specification noted by Janssen et al. ^11^. CellBender showed low cross-species removal under our tested settings but was the strongest of the three principal comparators for PT-marker leakage. Previous benchmarks reported favorable CellBender performance using different datasets and evaluation criteria ^4,11^. These differences reinforce the need to report removal and retention separately and to interpret rankings within the tested configurations.

scCDC targets gene-specific contamination through cluster-level subtraction ^6^. Its results are reported separately, including unsuccessful runs and their failure modes (Supplementary Table 2). Broader comparisons of configuration sensitivity remain necessary before generalizing the rankings observed here.

Several limitations are important for interpretation. The operational dose is not a calibrated contamination fraction, and ubiquitously expressed genes can inflate it (Supplementary Note 1). The released allocator uses nominal budgets to guide removal but does not enforce a hard per-cell cap; count monotonicity alone does not ensure preservation of endogenous RNA. In the high-ambient simulation, AmbiDose had lower clustering agreement than the uncorrected matrix, with errors concentrated in low-count, highly contaminated cells. Ambient and endogenous molecules remain difficult to distinguish when they contribute to the same gene–cell entries.

Protection also depends on coarse groups inferred from contaminated expression data or supplied by the user. Mixed groups, such as the pancreas cluster with both acinar and lymphoid expression, complicate interpretation. Mitochondrial genes retain a name-based protection rule, so the complete method is not gene-name independent. AmbiDose requires empty-droplet information, and debris-rich nucleus libraries may require additional modeling ^19,20^. Cross-species signal may include index hopping ^21^, which was not separately estimated. Finally, most benchmark datasets contributed to method development; only the GEM-X mixture and dorsal root ganglion library were held out.

These results support AmbiDose as an approach for reducing identifiable ambient signal while using cell-type structure to limit loss of lineage-enriched expression. Its strongest evidence in the present evaluation comes from the held-out species mixture and the marker-removal versus retention trade-off in high-background kidney nucleus libraries.

## Methods

AmbiDose is a Python package for ambient-RNA correction of droplet count matrices. It integrates with Scanpy and AnnData workflows. Analyses in this paper used version 0.5.11. Correction is performed through ambidose.denoise().

## Model

For cell c in sample s, let y_cg_ denote the observed count for gene g and n_c_ = Σg y_cg_ the total UMI count. The ambient profile χ_s_ is a probability vector with Σg χ_sg_ = 1. The operational dose ρ_c_ lies in [0, 1] and defines a nominal count-removal budget d_c_ = ρ_c_ n_c_. The term nominal distinguishes this allocation parameter from both a calibrated estimate of ambient molecules and a strict upper bound on realized removal.

AmbiDose estimates the ambient profile, resolves coarse cell groups, estimates and selects a cell-specific dose, and allocates removal using gene-level evidence and protection. The schematic expression 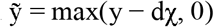 describes untyped profile-directed subtraction; the default typed path additionally uses pooled gene targets, protection, reallocation and clearance of genes classified as unexpressed and unprotected. In the default integer-count output, every entry satisfies 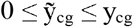. This entrywise constraint does not bound total removal by d_c_ and does not identify the source of removed molecules.

### Ambient profile from empty droplets

When a filtered barcode list is present, empty droplets are the complementary barcodes in a low-UMI empty band. Barcodes at or below the lower bound of that band are labeled other and are not used for χ. For each sample s, the ambient profile is the pooled empty-droplet composition

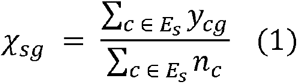

Ambient profiles are estimated separately for each sample. Stored profiles must have gene labels matching the count matrix; mismatches raise an error. At least 10 usable empty droplets are required. Empty-droplet data also provide the mean UMI count λ_e_, an empty-cloud UMI knee and a pooled NB2 overdispersion estimate φ for downstream diagnostics and background-dependent rules.

### Coarse cell types

Users may supply coarse cell-group labels. Otherwise, AmbiDose uses Leiden clustering ^22^ after Scanpy preprocessing ^23–25^. Groups with fewer than 10 cells do not undergo type-level clearance of unexpressed, unprotected genes and instead use the protected profile-directed path. Groups define evidence and protection sets; they do not replace cell-specific dose estimates with a group constant.

### Evidence pool and per-cell dose

Within each coarse type t of N_t_ cells, a gene is eligible for the ambient evidence pool U_t_ if its type mean is compatible with a pure-ambient ceiling. The default margin is Poisson:

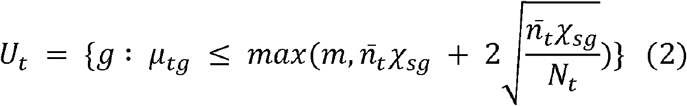

Here, μ_tg_ is the mean count of gene g in type t, and n □_t_ is the mean total UMI count. With the default floor m = 0, the ceiling identifies genes compatible with ambient-only expression; it does not establish that their counts are exclusively ambient. Genes classified as endogenous in every usable type are excluded, including genes above the ceiling in every type and genes satisfying the high-relative-expression collision criterion (r_t_ > 0.85 and mean > 0.003 × n _t_ in every type). Candidates are ranked by χ_sg_ and selected using a Lorenz-curve knee, subject to the minimum gene count of 8. When fewer than 8 candidates are available, a quantile-based estimate is used on the selected genes; its quantile is calibrated using empty droplets, with a fallback of 0.5 when calibration is unavailable. Ownership is not applied as a filter to this evidence pool.

For a type with sufficient candidate genes, the initial score is the Poisson maximum-likelihood estimate on the selected genes, including zero counts ^26^:

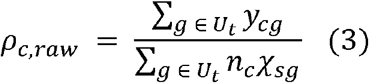

For samples with multiple usable types, finite initial estimates are shrunk on the log scale toward the sample median. Let θ_c_ = log(ρ_c,raw_ + 10^−8^), μ = median(θ), e_c_ = n_c_ Σg□U_t_ χ_sg_ and k be the median positive evidence exposure, or 8 if unavailable. The weight w_c_ = e_c_ / (e_c_ + k) gives ρ_c,fixed_ = clip(exp(w_c_ θ_c_ + (1 − w_c_)μ), 0, 1). This shrinks cells with limited evidence more strongly toward the sample median. In the exposure-based implementation, quantile-fallback estimates do not receive MLE evidence exposure. Cells without a finite initial estimate use the sample log-median when available. A sample without any finite initial estimates receives dose 0; samples with fewer than two usable types follow the untyped quantile path without this shrinkage step.

A two-component endogenous-plus-ambient mixture is also fitted with χ fixed and an updating endogenous profile. For cells with n_c_ > 0, the mixture estimate is selected when 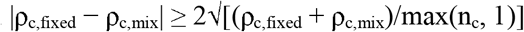 and the mixture status is fitted_converged or quantile_fallback. Otherwise, the fixed estimate is retained. The selected estimate is ρ_c_, and d_c_ = ρ_c_ n_c_ is passed to subtraction. Samples with fewer than two usable types use the untyped quantile estimate rather than the two-component fit. The per-sample diagnostic q = median(ρ□)n □/λ_e_ is also recorded; missing required empty-droplet information raises an error.

### Ownership and protection

Highly expressed endogenous genes can dominate the ambient profile and still satisfy the ceiling in equation (2). AmbiDose therefore assigns protection from differences in type-level mean expression. We use ownership to denote this algorithmic protection assignment, not exclusive biological expression.

For each gene, type means are sorted descending. Exclusive ownership compares each candidate’s mean, minus one sampling-error unit, with the runner-up. A tie leaves the gene unowned. Every type at or above that rank shares ownership if its mean exceeds the ownership floor. With the default floor, that threshold is 1 / median{N_t_ : N_t_ ≥ 10} inside the sample; a user-supplied max_type_mean replaces it. A merged fragment inherits its parent group’s ownership only when it is the sole member of that group, or when its own mean still exceeds every other parent group’s mean. For genes in the Lorenz-knee prefix of χ, ownership is restricted to a single winning type. Ownership is then stripped from any type whose r_t_ = mean / (n □χ) ≤ 1 if another type has r_t_ > 1.

Ownership affects subtraction: assigned genes are excluded from unexpressed-gene clearance and receive reduced profile-directed removal. It does not filter the dose evidence pool. Ribosomal and histone genes have no separate name-based exceptions. Mitochondrial symbols matching (^|_)mt[-_] receive a name-based protection mask; symbols such as MTOR are excluded. Before clipping to observed counts, native confidence q_g_ scales the profile-directed removal amount by (1 − 0.9q_g_); q_g_ = 1 therefore leaves 10% of the unweighted amount. Background-dependent rules can modify this protection (below). The rule does not guarantee a bound on endogenous UMI loss.

The gap-cascade rule leaves unprotected any gene that is ambiguously mid-level across several types with no fold gap. A second test targets that case. For each type t, an anchor gene set is built from genes with a confident owner in some other type (mean at least 3-fold above the runner-up), and an independent per-type ambient-dose estimate ρ_t,anchor_ is computed from that anchor set alone. A candidate gene g (unexpressed by equation (2) in type t) is additionally protected if its type mean exceeds 2.5 times the anchor-implied ambient expectation n _t_ χg ρ_t,anchor_, and its log-CP10K expression across type t’s cells has adjusted R^2^ ≥ 0.01 against latent cell-state components (truncated SVD, 20 components) fit only on the confident-owner reference genes. Extra protection is skipped for a type whose own ρ_t,anchor_ falls below 0.1. These thresholds are fixed for every dataset. The anchor changes protection only; it does not change the dose.

### Count allocation and its guarantees

For sample s and type t, the sum of nominal cell budgets is D_t_ = Σc□t d_c_. Initial gene-level profile-directed targets are limited by the observed type-level count and D_t_ χ_sg_, with native-confidence weighting applied before subsequent reallocation. Unprotected targets are modestly reweighted using the nonnegative within-type correlation between normalized gene abundance and dose fraction, preserving target mass when feasible. Unsaturated unprotected targets are allocated through dose-weighted per-cell targets; saturated targets and protected-gene targets are allocated gene by gene across cells. Protected-gene allocation uses a reliability-weighted blend of dose ranks and anchor-based ambient-rate ranks. Each allocation is limited by the observed count available at that entry.

Unused profile-directed target mass can be reallocated within the same sample and type to genes outside the additional-clearance set. Eligible genes satisfy r_t_ = mean/(n□ χ) ≤ 1. Nonnegative slopes of normalized gene abundance against cell-specific ρ provide ambient-response support. This support restricts both allocation weight and capacity; for protected genes it is further reduced by (1 − q_g_). Unsupported capacity is left unused.

After profile-directed allocation, the default additional-clearance step calculates each cell’s nonnegative residual allowance as max(d_c_ − R_c,rank1_, 0). The sum of these allowances scales the pooled targets for genes classified as unexpressed and unprotected, and the allowances are used as cell weights during gene-wise allocation. The allocator does not enforce each allowance as a joint cap across genes. Integer targets are rounded half up, limited by available counts and distributed by largest fractional remainder with deterministic identifier-based tie breaking. No final row-wise projection back to d_c_ is performed. Consequently, the released implementation guarantees integer output and 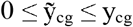, but not 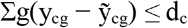. Realized removal and the ratio of realized removal to d_c_ are recorded for quality control. When unexpressed-gene mass or estimated ambient mass is consistent with the empty-droplet background, additional clearance and leftover reallocation are skipped and the initial profile-directed target is reduced. A separate low-background rule reduces native protection and reweights χ toward unexpressed or low-confidence genes when the estimated ambient mass per cell is above 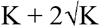 but no greater than 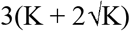, where K is the empty-cloud UMI knee, provided that the empty-consistency rule has not already applied.

### Implementation

AmbiDose integrates with Scanpy and AnnData workflows. The reported analyses used the same AmbiDose configuration across datasets. Parameters remain user-configurable.

By default, denoise() treats a Cell Ranger filtered barcode list ^27^ as cells as-is. That list is not re-called. EmptyDrops ^28^, DIEM ^19^ or DropletQC ^29^, or cell_calling=‘chi’ are used only when no filtered list is supplied, or when that list is known to be over-called. Doublet flagging with scanpy Scrublet ^30^ is optional and is not part of the default path used in this paper. Empty-band defaults are empty_umi_min = 0 and empty_umi_max = 100. At least min_empty = 10 usable empty droplets are required.

When users do not supply type_key, Leiden resolution is selected by scFair when installed ^31^: 0.08 if fewer than 12 estimated populations, 0.2 if 12–19, and 0.35 if 20 or more. scFair does not name cell types. If it is unavailable or errors, the resolution is 0.35. The comparisons in this paper used that fallback. On libraries with at least 20,000 cells, typing_fast=True (the default) builds a cheaper neighborhood graph. Groups with fewer than MIN_TYPE_CELLS = 10 cells skip type-level soupOnly clearance.

Evidence-pool defaults include NOISE_K = 2.0, max_type_mean = None (m = 0), U_MAX_LIBRARY_FRAC = 0.003 and MIN_GENES = 8. Dose estimation uses evidence_mode=“exposure” and SHRINK_K = 8 when the evidence median is unavailable. Ownership uses OWNER_GAP_SE_Z = 1. Default subtraction uses empirical_margin=True and cross_type_anchor=True. The parameters cap_high_u_to_remaining=True and high_u_remaining_multiplier=1.0 scale pooled additional-clearance targets using summed nonnegative residual allowances; they do not impose a hard per-cell removal cap.

The returned AnnData stores the integer corrected matrix in layers[“ambidose_denoised”] and X, and preserves input counts in layers[“raw_counts”]. The field obs[“ambidose_rho_trust”] contains diagnostic labels: ok, low_evidence, ceiling_risk, type_structure_risk, under_execution, over_removal and not_cell. These labels are not calibrated probabilities of correctness. For positive d_c_, under_execution indicates realized removal below half the nominal budget, and over_removal flags an execution ratio above 1.05. The latter also flags removal exceeding half of a cell’s raw UMIs. These diagnostics report the relationship between estimated dose and executed removal rather than a formal guarantee of endogenous-signal preservation.

### Datasets and comparators

Evaluation used two human–mouse mixtures (hgmm12k ^16^ and the 10x/SoupX Mixture dataset ^1^), a 24-sample kidney-tumor cohort ^14^, fetal liver ^15^, GSE147203 ^10^, GSE218853 ^11^, PBMC4k ^32^ for contamination injection, GSE125588 ^33^ for the pancreas case study, and GSE265817 for the held-out dorsal-root ganglion library ^18^. The plate-based negative control used E-MTAB-5061 ^34^. The held-out GEM-X validation used the public 10x Genomics 10k 1:1 human HEK293T–mouse NIH3T3 Chromium GEM-X 3′ dataset (10k-hgmm-3p-gemx; Cell Ranger 8.0.0) ^35^. Analysis used the downloaded raw matrix (72,302 features × 2,095,329 barcodes) and the matched filtered HDF5 (12,735 barcodes); those files were obtained after the AmbiDose protocol was locked and were not used for parameter or feature selection. Comparisons used the intersection of retained cell barcodes when methods differed in cell calling; a match below 90% of AmbiDose’s own cell count was treated as a failed comparison and excluded, not silently padded. Software version 0.5.11 is the released estimator described in this paper.

Comparator settings were not adjusted after examining a library’s leak ratio, cross-species removal or hemoglobin retention. AmbiDose used one configuration across datasets, but its parameters were selected on the development collection. The GEM-X and dorsal root ganglion datasets were held out. These comparisons therefore evaluate the stated configurations rather than the best achievable performance of every method.

CellBender 0.3.2 used the documented defaults (--fpr 0.01, --epochs 150, --estimator mckp) on every dataset with a CellBender row. expected-cells and total-droplets-included were set from each library’s Cell Ranger cell count so training would initialize. They were not swept to improve a reported metric. SoupX 1.6.2 was run as published: autoEstCont received a correction-free frozen Seurat v5 cluster file ^36^, the soup profile used soupRange = (0, 100) when empty droplets were available, autoEstCont used its default prior, priorRho = 0.05, and estimated the contamination fraction rather than fixing it at 0.05, and adjustCounts ran at cell level. DecontX is celda 1.28.0.

The figure runs are decontX(z = NULL) with no background matrix. The function accepts empty droplets through background; that rerun is Supplementary Table 7. FastCAR is cited for mechanism and was not run as official software in the tables. scCDC 1.4 was run at documented defaults; run success and failure modes on the kidney-tumor and fetal-liver cohorts are reported in Supplementary Table 2. R session: R 4.6.1. Comparator versions and configurations are summarized in Supplementary Table 3.

### Evaluation metrics

For species mixtures, cross-species UMIs in confidently assigned singlets were used as a contamination proxy. Sensitivity is the fraction of cross-species UMIs removed; specificity is the fraction of same-species UMIs retained; precision is the number of removed cross-species UMIs divided by all removed UMIs. Same-species counts include same-species ambient RNA and are not equivalent to endogenous counts. Supplementary Fig. 1 uses each species’ central 20% of library size and excludes barcodes with more than 10% cross-species UMIs. For primary tissues, on-target expression is the mean expression of a type’s canonical markers in that type, and off-target expression is the mean expression of other types’ markers. The leak ratio is off-target divided by on-target expression. On-target UMI retention is the corrected-to-raw ratio for the marker panel. Count inflation is the number of corrected entries exceeding input counts. Variance decomposition used η^2^ = SSbetween/SStotal with cells as observational units and estimated dose as the response. Groupings included coarse type, Leiden cluster, sample and their within-sample combinations. Hemoglobin retention is the corrected-to-raw ratio of mean UMI counts for HBA1, HBA2 and HBB in erythroid or non-erythroid cells, using matched barcodes and raw-derived coarse labels.

For GSE218853, gene–cell positions were classified using the authors’ CAST-strain genotype masks. Foreign-positive positions had nonzero foreign-allele fractions, and foreign-negative positions had zero foreign-allele fractions. Positions without informative SNP coverage or with missing allele calls were excluded. Mean removal at foreign-positive positions was calculated as the per-gene mean of min(1, |raw − corrected|/raw); retention at foreign-negative positions was the per-gene mean of max(0, 1 − |raw − corrected|/raw). These position-level scores differ from molecule-origin ARS and ERS bounds in the simulations. The removed-UMI share is the number removed from foreign-positive positions divided by the number removed from both scored compartments. A foreign-positive position can contain both foreign and endogenous molecules, and a zero observed foreign-allele fraction does not establish absence of ambient RNA.

The genotype reference is the UMI-weighted mean (Σ n_c_ρ _c_ / Σ n_c_) of the authors’ per-cell maximum-likelihood background estimate. This estimate uses a binomial model over informative SNP loci (method of Heaton et al. ^37^, as implemented by Janssen et al. ^11^) that extrapolates from cross-genotype signal to a whole-cell estimate, with a profile-likelihood 95% confidence interval per cell. It is defined only on genotype-matched CAST cells, a minority of each library (2–30% of cells, lowest on nuc2 and nuc3); because this population differs from the full library, we do not compare library-wide removal totals with it. Mean informative-SNP coverage per cell is 564– 816 for rep1–rep3 and nuc3 and 127 for nuc2, the low-coverage library reported by the authors.

Fig. 5b,c score the 75 genes of the 83-gene PanglaoDB ^17^ proximal-tubule panel that are present in the raw 10x gene index. The same 75 genes are used for every method, and the panel was not derived from this dataset’s expression ranking. We report PT-marker leakage (mean detection fraction in non-PT cells), specificity (CP10K-normalized log2 fold change, PT versus other types, pseudocount 1) and PT-internal signal lost (1 minus the corrected/raw UMI ratio for these genes within PT cells, with the corresponding within-PT detection-rate drop). Leakage and PT-internal signal lost are read together so that leakage reduction is weighed against on-target cost.

Synthetic contamination was injected into PBMC4k ^32^ cells and empty droplets using INS, GCG, CPA1, ALB, MYH6 and TNNT2, which were absent from the clean background. Cell-specific ambient fractions were drawn from a clipped normal distribution and counts from a negative-binomial model. Recovery denotes the fraction of injected UMIs removed. For Supplementary Fig. 2, d_c_ was compared with the intended ambient count, defined as library size multiplied by the simulated fraction; the reported slope is from ordinary least squares. Library-size windows are quintiles. Dose-median and within-group shuffle ablations keep χ, ownership and the subtraction procedure fixed; the shuffle uses seed 0. The single-group ablation assigns all cells the same label before dose estimation and subtraction. Denoising uses inferred Leiden groups rather than true cell-line labels. ARI compares independently inferred clusters with cell-line identity. For the cell-line simulations, ARS and ERS are bounded because the corrected matrix does not label the molecular source of removal. At an entry with ambient count A, endogenous count E and removal R, ambient removal lies between max(0, R − E) and min(R, A). Summing these bounds and dividing by total injected ambient counts gives the ARS bounds. Endogenous removal lies between max(0, R − A) and min(R, E); subtracting the corresponding normalized upper and lower removal bounds from 1 gives the lower and upper ERS bounds, respectively. These are attribution bounds, not statistical confidence intervals.

### Statistical analysis and reproducibility

Paired differences in leak ratio were evaluated with two-sided Wilcoxon signed-rank tests. To account for libraries sharing a donor, we report comparisons after averaging libraries within donor (7 kidney donors and 14 fetal-liver donors). Donor-level P values are reported without multiplicity adjustment. Library-level analyses used 23 evaluable kidney libraries and 39 fetal-liver libraries; one kidney library with only one scored type was excluded. A Bonferroni threshold of 0.0083 was used for the six library-level comparisons, which do not account for within-donor dependence. Paired bootstrap intervals for mean library-level differences used 10,000 resamples with replacement and a fixed seed. For the two original species-mixture libraries, cell-level bootstrap intervals used 10,000 resamples stratified by species and characterize within-experiment sampling uncertainty rather than biological-replicate variability. Count-removal and retention metrics used raw counts; marker-specific normalization is described above. Software tests evaluate count monotonicity and sparse-matrix behavior.

AmbiDose is CPU-only and does not require a GPU. Runtime was measured in a controlled GEM-X benchmark on the same 12,735 filtered barcodes used for the held-out mixture comparison, with AmbiDose, SoupX and DecontX each run on four CPU cores and CellBender on GPU with eight CPU threads (GNU time -v; Supplementary Table 4). Reported speed ratios are specific to that benchmark and hardware, not universal software claims.

## Data availability

No new sequencing data were generated for this study. hgmm12k is the public 10x Genomics 12k 1:1 HEK293T– NIH3T3 mixture ^16^. PBMC4k is the public 10x Genomics 4k PBMC library ^32^. The 10k 1:1 human HEK293T– mouse NIH3T3 Chromium GEM-X 3′ mixture is the public 10x dataset 10k-hgmm-3p-gemx (Cell Ranger 8.0.0 filtered HDF5, 12,735 barcodes) ^35^. The Mixture dataset is the data released with SoupX ^1^. GSE147203 ^10^, GSE218853 ^11^, GSE125588 ^33^ and the uninjured mouse DRG library GSE265817 (GSM8229618) ^18^ are available from GEO. E-MTAB-5061 is available from ArrayExpress ^34^. Kidney-tumor libraries used here are public 10x count matrices listed in Supplementary Table 5; sample identifiers match the Young laboratory collection ^14^.

Fetal-liver libraries are public 10x count matrices listed in Supplementary Table 6; FCAImmP and 4834STDY identifiers match the Human Cell Atlas fetal-liver atlas ^15^.

## Code availability

The reported analyses used AmbiDose version 0.5.11. Source code is available at https://github.com/leelieber2025/AmbiDose under the Apache License 2.0. The version 0.5.11 record is https://doi.org/10.5281/zenodo.22882960 (concept DOI, https://doi.org/10.5281/zenodo.22278199). The package can be installed with pip install ambidose==0.5.11. Documentation is available at https://ambidose.readthedocs.io.

## Acknowledgements

Analyses use publicly available de-identified datasets; the original studies obtained the relevant approvals. The authors used personal computational resources.

## Author contributions

Z.L. conceived the method, implemented the software, designed and performed the analyses, and wrote the manuscript. A.W.J. edited the manuscript. S.L. implemented the software.

