## Supplemental for "AmbiDose: per-cell removal budgets for ambient RNA contamination"

**Supplementary information**

**Supplementary Note 1 Bias from ubiquitously expressed genes**

A controlled example illustrates bias in the operational dose when ubiquitously expressed genes enter the evidence pool. Eighty shared genes each had a mean count of 0.4, approximately 0.2% of the library and below the 0.3% collision threshold (U_MAX_LIBRARY_FRAC = 0.003). The χ-ranked evidence pool contained 76–80 of these genes among 100 candidates. With fixed settings, median ρ was 0.218, 0.279 and 0.336 for true scales of 0.02, 0.10 and 0.20, respectively. This pattern indicates an approximately additive offset rather than a constant multiplicative bias. In primary tissues, the distribution of mean/(n̄χ) among uniform genes did not clearly separate ambient and housekeeping expression: both modes lay near or above the ceiling. Alternative rankings based on empty-versus-cell enrichment, low ratios or low detection either reduced estimated doses toward zero or inflated species-mixture estimates. A two-component model of gene-level count-to-exposure ratios recovered the controlled example but underestimated dose in kidney tumor and fetal liver by excluding genes carrying modest contamination. These results motivate interpreting ρ as an operational score rather than a calibrated contamination fraction. Entrywise count monotonicity does not resolve this estimation bias or guarantee endogenous-molecule retention.

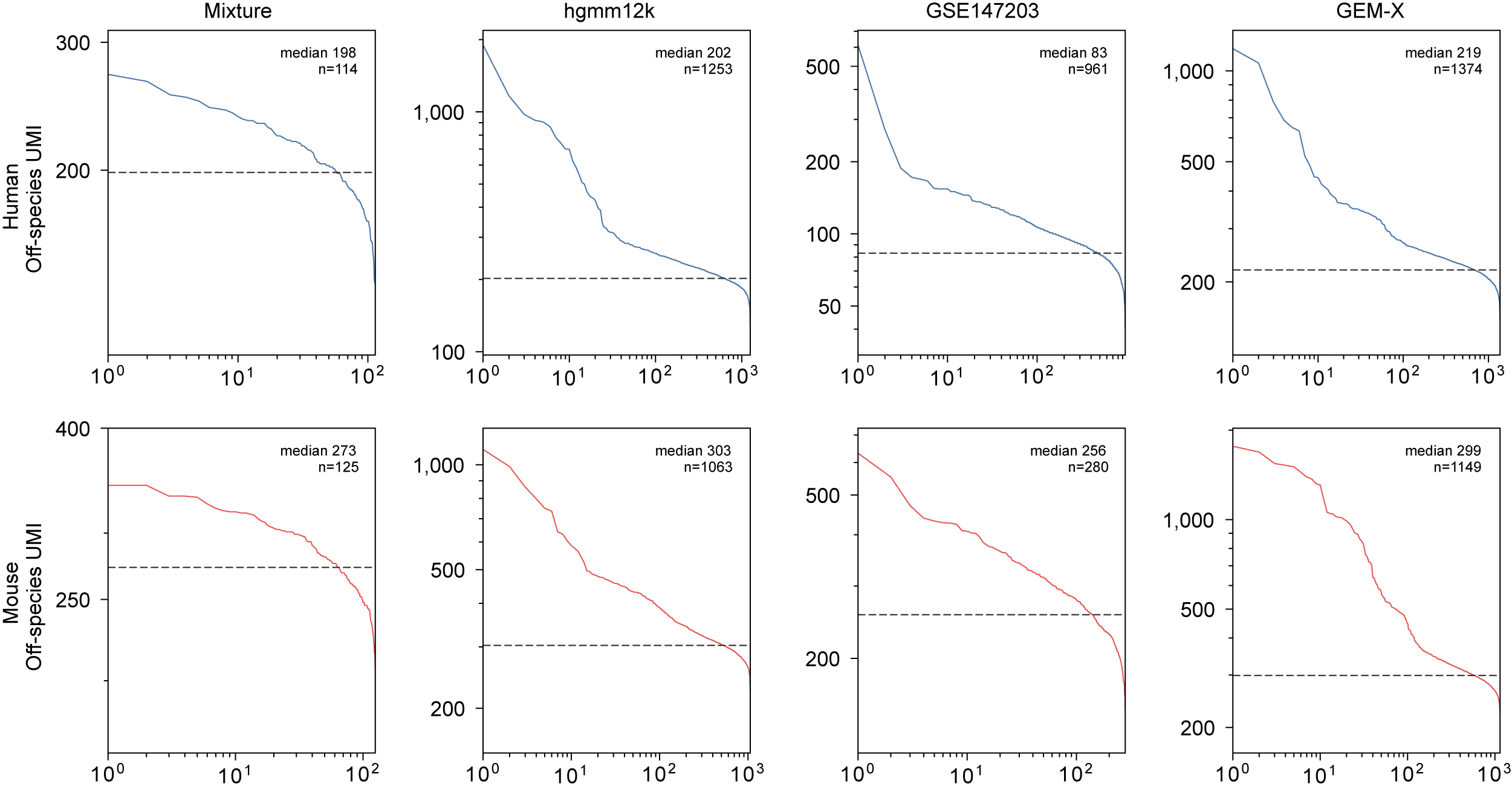

**Supplementary Fig. 1 | Off-species UMI counts on four species-mixing libraries, ordered from high to low.** Columns are Mixture, hgmm12k, GSE147203 and GEM-X. Top row, human cells; bottom row, mouse cells. Each panel uses only cells in that species’ central 20% of library size, to reduce differences attributable to sequencing depth or species. Barcodes with off-species UMIs above 10% of their total are omitted. The dashed line is the median. The printed n is the number of cells in that panel.

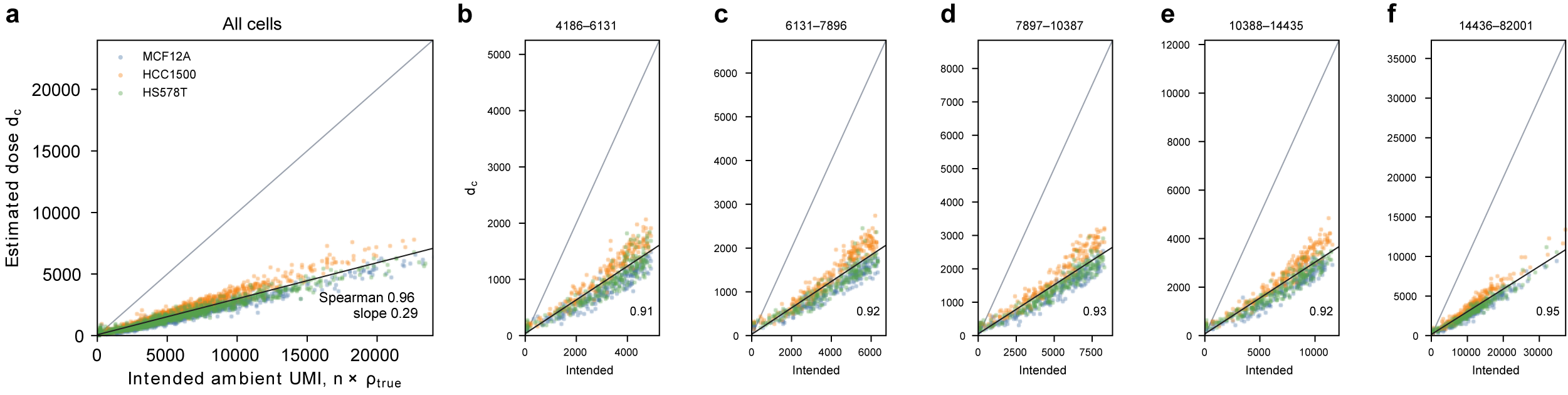

**Supplementary Fig. 2 | Nominal removal budget against the intended ambient UMI count on the Figure 4 simulation.** Denoising used Leiden groups, not the true cell lines. (a) Each point is a cell, colored by the true line. The gray line is y = x. The black line is ordinary least squares of d_c on n_c × ρ_true (Spearman 0.96; slope 0.29). (b–f) The same cells split into five library-size quintiles; the printed number is the Spearman correlation inside that window.

**Supplementary Table 1 | Top 20 genes by ambient weight χ in GSE125588 (mouse pancreas). Cumulative mass is the running share of total χ. The panel is dominated by acinar-associated transcripts and also includes Hbb-bs and Ins2.**

| **Rank** | **Gene** | **χ weight (%)** | **Cumulative χ mass (%)** |
| --- | --- | --- | --- |
| 1 | Clps | 10.94 | 10.94 |
| 2 | Ctrb1 | 9.42 | 20.36 |
| 3 | Cela2a | 6.85 | 27.21 |
| 4 | Prss2 | 6.65 | 33.86 |
| 5 | Cela1 | 4.15 | 38.02 |
| 6 | Prss3b | 3.92 | 41.94 |
| 7 | Try5 | 3.45 | 45.39 |
| 8 | Cela3b | 3.36 | 48.75 |
| 9 | Try4 | 3.02 | 51.77 |
| 10 | Sycn | 2.53 | 54.30 |
| 11 | Pnlip | 2.20 | 56.50 |
| 12 | Reg1 | 2.05 | 58.55 |
| 13 | Rnase1 | 1.46 | 60.01 |
| 14 | Cpb1 | 1.38 | 61.39 |
| 15 | Zg16 | 1.28 | 62.68 |
| 16 | Ctrl | 1.13 | 63.81 |
| 17 | Cpa1 | 1.08 | 64.89 |
| 18 | Hbb-bs | 0.83 | 65.72 |
| 19 | Pnliprp1 | 0.79 | 66.52 |
| 20 | Ins2 | 0.59 | 67.11 |

**Supplementary Table 2 | scCDC comparison (documented defaults). For GSE218853 rows, FA+ / FA− are position-level foreign-allele-positive removal and foreign-allele-negative retention (Methods), not the molecule-level ARS/ERS of Fig. 4.**

| **Dataset** | **Metric** | **scCDC** | **AmbiDose** |
| --- | --- | --- | --- |
| GEM-X | Sensitivity / Specificity / Precision | 0.135 / 0.977 / 0.063 | 0.920 / 0.996 / 0.730 |
| GSE218853 rep1 | FA+ removal / FA− retention | 0.0003 / 0.9998 | 0.031 / 0.981 |
| GSE218853 rep2 | FA+ removal / FA− retention | 0.0025 / 0.9988 | 0.034 / 0.989 |
| GSE218853 rep3 | FA+ removal / FA− retention | 0.000 / 1.000 | 0.017 / 0.996 |
| GSE218853 nuc2 | FA+ removal / FA− retention | 0.017 / 0.989 | 0.293 / 0.882 |
| GSE218853 nuc3 | FA+ removal / FA− retention | 0.000 / 1.000 | 0.201 / 0.904 |

scCDC 1.4, documented defaults, not retuned per dataset (same policy as SoupX/DecontX/CellBender). On the kidney-tumor and fetal-liver cohorts, scCDC succeeded on 16/24 and 35/39 libraries respectively; failures (8 kidney, 4 fetal-liver) were due to a smooth.spline numerical error in scCDC’s own threshold-fitting step (7 libraries) or no cluster meeting scCDC’s own min.cell=100 requirement (5 libraries). Successful runs removed a median of 10.9% (kidney) and 3.2% (fetal liver) of UMIs per library; no leak-ratio-equivalent ground truth was computed for scCDC on these cohorts, so these are reported as UMI-removal context only, not an accuracy comparison.

**Supplementary Table 3 | Comparator configuration**

| **Method** | **Version** | **Input** | **Key parameters** | **GPU** | **Cell calling / barcodes** |
| --- | --- | --- | --- | --- | --- |
| AmbiDose | 0.5.11 | Cell Ranger raw h5/mtx + filtered barcodes | denoise() defaults in 0.5.11: evidence_mode="exposure", cross_type_anchor=True, empirical_margin=True, cap_high_u_to_remaining=True, NOISE_K=2.0, MIN_TYPE_CELLS=10, U_MAX_LIBRARY_FRAC=0.003, empty band 0 < n_UMI ≤ 100; identical across datasets | No | Cell Ranger filtered barcodes as default input path; reference set |
| CellBender | 0.3.2 | Unfiltered droplet matrix | --fpr 0.01; --epochs 150; --estimator mckp (tool defaults) | Yes | Own posterior; intersected with AmbiDose barcodes |
| SoupX (R) | 1.6.2 | Filtered toc + raw tod where available | autoEstCont on correction-free Seurat v5 clusters; priorRho 0.05; adjustCounts cell-level; soupRange (0, 100) | No | Same barcodes as AmbiDose |
| DecontX (R) | celda 1.28.0 | Filtered toc | decontX(z = NULL), package defaults, no labels | No | Same barcodes as AmbiDose |

**Supplementary Table 4 | Computational resource benchmark**

Runtimes are for one library, the 10x GEM-X mixture, using its 12,735 filtered barcodes. CellBender used a GPU; the other three used four CPU cores.

| **Method** | **Version** | **Hardware** | **Representative run** | **Runtime** | **Peak memory** | **Measurement basis** |
| --- | --- | --- | --- | --- | --- | --- |
| SoupX | 1.6.2 | CPU; 4 cores | GEM-X; 12,735 cells | 13 min 50 s | 14.04 GiB RSS | GNU time -v; controlled run |
| DecontX | celda 1.28.0 | CPU; 4 cores | GEM-X; 12,735 cells | 14 min 46 s | 7.94 GiB RSS | GNU time -v; controlled run |
| CellBender | 0.3.2 | GPU (CUDA); 8 CPU threads | GEM-X; MCKP, 150 epochs | ~8 h 56 min | 19.99 GiB RSS; 7.55 GiB VRAM | Full runtime from strict log; RSS from checkpoint-resume GNU time -v |
| AmbiDose | 0.5.11 | CPU; 4 cores | GEM-X; 12,735 cells | 2 min 50 s | 10.66 GiB RSS | GNU time -v; controlled 4-core run |

**Supplementary Table 5 | Kidney-tumor per-sample leak ratios**

Twenty-three of 24 kidney samples had at least two marker-scored types. Wilms2_Kid_T_ldc_1_3 (236 cells, one scored type) is listed with dashes because a leak ratio needs an off-target type and an on-target type. Versus CellBender, AmbiDose had lower leak in 21 of 23 samples with 2 ties and 0 losses (Wilcoxon signed-rank *P* = 5.96 × 10^−5^; paired bootstrap 95% CI for AmbiDose minus CellBender, −0.0308 to −0.0034). Versus SoupX, 17 of 23 (ties 2; *P* = 0.0087; CI −0.0212 to −0.0004). Versus DecontX, 14 of 23 (ties 1; *P* = 0.071; CI −0.0152 to −0.0004).

| **Sample** | ***N* cells** | ***N* types** | **AmbiDose leak** | **SoupX leak** | **DecontX leak** | **CellBender leak** |
| --- | --- | --- | --- | --- | --- | --- |
| RCC1_Kid_T_ldc_1_1 | 214 | 6 | 0.0298 | 0.0304 | 0.0253 | 0.0327 |
| RCC1_Kid_T_ldc_1_2 | 208 | 7 | 0.0183 | 0.0196 | 0.0142 | 0.0222 |
| RCC1_Kid_T_ldc_2_1 | 1061 | 7 | 0.0017 | 0.0018 | 0.0014 | 0.0018 |
| RCC1_Kid_T_ldc_2_2 | 1050 | 8 | 0.0020 | 0.0020 | 0.0031 | 0.0020 |
| RCC2_Kid_T_ldc_1_1 | 1312 | 8 | 0.0266 | 0.0309 | 0.0288 | 0.0347 |
| RCC2_Kid_T_ldc_1_2 | 1595 | 8 | 0.0047 | 0.0057 | 0.0039 | 0.0061 |
| RCC2_Kid_T_ldc_2_1 | 2675 | 8 | 0.0057 | 0.0060 | 0.0054 | 0.0057 |
| RCC2_Kid_T_ldc_2_2 | 2674 | 8 | 0.0049 | 0.0055 | 0.0055 | 0.0059 |
| VHL_Kid_T_ldc_1_1 | 989 | 7 | 0.0174 | 0.0189 | 0.0172 | 0.0189 |
| VHL_Kid_T_ldc_1_2 | 875 | 6 | 0.0303 | 0.0322 | 0.0287 | 0.0337 |
| Wilms1_Kid_R_ldc_1_1 | 149 | 6 | 0.0362 | 0.0435 | 0.0411 | 0.0482 |
| Wilms1_Kid_R_ldc_1_2 | 139 | 7 | 0.0494 | 0.0537 | 0.0671 | 0.0802 |
| Wilms1_Kid_R_ldc_1_3 | 159 | 7 | 0.0272 | 0.0269 | 0.0277 | 0.0338 |
| Wilms1_Kid_T_ldc_1_1 | 53 | 3 | 0.0549 | 0.0451 | 0.0586 | 0.0585 |
| Wilms1_Kid_T_ldc_1_2 | 153 | 6 | 0.0162 | 0.0180 | 0.0180 | 0.0194 |
| Wilms1_Kid_T_ldc_1_3 | 88 | 5 | 0.0286 | 0.0218 | 0.0311 | 0.0290 |
| Wilms2_Kid_T_ldc_1_1 | 266 | 2 | 0.0137 | 0.1137 | 0.0815 | 0.1429 |
| Wilms2_Kid_T_ldc_1_2 | 297 | 2 | 0.0175 | 0.1024 | 0.0781 | 0.1337 |
| Wilms2_Kid_T_ldc_1_3 | 236 | 1 | — | — | — | — |
| Wilms3_Kid_T_ldc_1_1 | 213 | 7 | 0.0325 | 0.0369 | 0.0309 | 0.0392 |
| Wilms3_Kid_T_ldc_1_2 | 253 | 6 | 0.0268 | 0.0301 | 0.0284 | 0.0317 |
| Wilms3_Kid_T_ldc_1_3 | 230 | 7 | 0.0273 | 0.0324 | 0.0299 | 0.0345 |
| pRCC_Kid_T_ldc_1_1 | 739 | 7 | 0.0031 | 0.0031 | 0.0032 | 0.0033 |
| pRCC_Kid_T_ldc_1_2 | 654 | 5 | 0.0026 | 0.0025 | 0.0026 | 0.0029 |

**Supplementary Table 6 | Fetal-liver per-sample leak ratios**

AmbiDose had lower leak than CellBender in 38 of 39 samples (Wilcoxon *P* = 7.28 × 10^−12^; paired bootstrap 95% CI −0.0276 to −0.0155). Versus SoupX, 29 of 39 (ties 1; *P* = 1.41 × 10^−4^; CI −0.0146 to −0.0004). Versus DecontX, 31 of 39 (ties 1; *P* = 8.22 × 10^−6^; CI −0.0332 to −0.0106).

| **Sample** | ***N* cells** | ***N* types** | **AmbiDose leak** | **SoupX leak** | **DecontX leak** | **CellBender leak** |
| --- | --- | --- | --- | --- | --- | --- |
| F16_liver_CD45neg_4834STDY7002878 | 1212 | 4 | 0.0148 | 0.0710 | 0.0523 | 0.0663 |
| F16_liver_CD45pos_4834STDY7002877 | 156 | 4 | 0.0073 | 0.0271 | 0.0228 | 0.0310 |
| F17_liver_CD45neg_4834STDY7038751 | 747 | 3 | 0.0259 | 0.0204 | 0.1059 | 0.0368 |
| F17_liver_CD45pos_4834STDY7038750 | 287 | 4 | 0.0091 | 0.0122 | 0.0112 | 0.0107 |
| F17_liver_Total_4834STDY7002882 | 1247 | 8 | 0.0163 | 0.0200 | 0.0123 | 0.0207 |
| F21_liver_CD45neg_FCAImmP7179364 | 4435 | 7 | 0.0303 | 0.0404 | 0.0497 | 0.0622 |
| F21_liver_CD45pos_FCAImmP7179363 | 3005 | 7 | 0.0145 | 0.0157 | 0.0190 | 0.0205 |
| F22_liver_CD45neg_FCAImmP7198629 | 3720 | 8 | 0.0154 | 0.0156 | 0.0211 | 0.0274 |
| F22_liver_CD45pos_FCAImmP7198628 | 2660 | 7 | 0.0153 | 0.0170 | 0.0357 | 0.0233 |
| F22_liver_Total_FCAImmP7198434 | 4823 | 4 | 0.0372 | 0.1234 | 0.1267 | 0.1009 |
| F23_liver_CD45neg_FCAImmP7198631 | 3152 | 7 | 0.0139 | 0.0118 | 0.0181 | 0.0250 |
| F23_liver_CD45pos_FCAImmP7198630 | 2982 | 7 | 0.0177 | 0.0219 | 0.0225 | 0.0275 |
| F29_liver_CD45neg_FCAImmP7277553 | 3529 | 8 | 0.0267 | 0.0233 | 0.0368 | 0.0430 |
| F29_liver_CD45pos_FCAImmP7277552 | 2410 | 7 | 0.0115 | 0.0120 | 0.0123 | 0.0156 |
| F30_liver_CD45neg_FCAImmP7277561 | 4513 | 8 | 0.0290 | 0.0289 | 0.0460 | 0.0525 |
| F30_liver_CD45pos_FCAImmP7277560 | 5758 | 8 | 0.0120 | 0.0103 | 0.0264 | 0.0119 |
| F32_liver_CD45neg_FCAImmP7316895 | 3411 | 6 | 0.0212 | 0.0260 | 0.0436 | 0.0587 |
| F32_liver_CD45pos_FCAImmP7316894 | 3649 | 8 | 0.0094 | 0.0119 | 0.0090 | 0.0156 |
| F33_liver_CD45neg_FCAImmP7316892 | 3287 | 6 | 0.0172 | 0.0485 | 0.0420 | 0.0730 |
| F33_liver_CD45neg_FCAImmP7316893 | 3275 | 6 | 0.0197 | 0.0391 | 0.0350 | 0.0625 |
| F33_liver_CD45pos_FCAImmP7316889 | 3672 | 8 | 0.0197 | 0.0152 | 0.0061 | 0.0234 |
| F33_liver_CD45pos_FCAImmP7316890 | 3820 | 6 | 0.0208 | 0.0219 | 0.0080 | 0.0269 |
| F33_liver_CD45pos_FCAImmP7316891 | 3551 | 6 | 0.0205 | 0.0214 | 0.0653 | 0.0301 |
| F34_liver_CD45neg_FCAImmP7352194 | 3265 | 6 | 0.0174 | 0.0238 | 0.0285 | 0.0502 |
| F34_liver_CD45neg_FCAImmP7352195 | 3269 | 7 | 0.0169 | 0.0220 | 0.0302 | 0.0522 |
| F34_liver_CD45pos_FCAImmP7352192 | 4368 | 8 | 0.0110 | 0.0123 | 0.0078 | 0.0178 |
| F34_liver_CD45pos_FCAImmP7352193 | 4609 | 8 | 0.0107 | 0.0135 | 0.0099 | 0.0172 |
| F35_liver_CD45neg_FCAImmP7462239 | 3026 | 7 | 0.0385 | 0.0513 | 0.0653 | 0.0972 |
| F35_liver_CD45pos_FCAImmP7462237 | 2790 | 8 | 0.0094 | 0.0167 | 0.0094 | 0.0210 |
| F38_liver_CD45neg_FCAImmP7528288 | 2559 | 6 | 0.0103 | 0.0382 | 0.0338 | 0.0759 |
| F38_liver_CD45neg_FCAImmP7528289 | 2465 | 4 | 0.0229 | 0.0340 | 0.0420 | 0.0706 |
| F38_liver_CD45pos_FCAImmP7528286 | 2179 | 6 | 0.0174 | 0.0229 | 0.2165 | 0.0292 |
| F38_liver_CD45pos_FCAImmP7528287 | 1900 | 6 | 0.0179 | 0.0227 | 0.0338 | 0.0300 |
| F41_liver_CD45neg_FCAImmP7555847 | 3598 | 5 | 0.0262 | 0.0261 | 0.0825 | 0.0528 |
| F41_liver_CD45pos_FCAImmP7555846 | 3117 | 7 | 0.0122 | 0.0103 | 0.0126 | 0.0149 |
| F45_liver_CD45neg_FCAImmP7579211 | 1877 | 7 | 0.0048 | 0.0034 | 0.0034 | 0.0098 |
| F45_liver_CD45neg_FCAImmP7579223 | 2012 | 6 | 0.0201 | 0.0361 | 0.0309 | 0.0548 |
| F45_liver_CD45pos_FCAImmP7579210 | 1832 | 8 | 0.0118 | 0.0118 | 0.0140 | 0.0133 |
| F45_liver_CD45pos_FCAImmP7579222 | 2141 | 7 | 0.0190 | 0.0305 | 0.0349 | 0.0477 |

Supplementary Table 7 | DecontX with empty-droplet background on four species-mixture libraries. Figure runs used decontX(z = NULL) and did not pass background. This rerun passes the same empty droplets used for chi, still with z = NULL. Same-species retention is not native-UMI retention. GEM-X is included.

| **Library** | **No background: sensitivity / specificity / precision** | **With empty droplets: sensitivity / specificity / precision** |
| --- | --- | --- |
| hgmm12k | 0.659 / 0.942 / 0.377 | 0.796 / 0.944 / 0.433 |
| Mixture | 0.507 / 0.996 / 0.616 | 0.874 / 0.986 / 0.448 |
| GSE147203 | 0.758 / 1.000 / 0.992 | 0.846 / 0.981 / 0.478 |
| GEM-X | 0.448 / 0.996 / 0.532 | 0.728 / 0.976 / 0.407 |
